# Discovery of directional chromatin-associated regulatory motifs affecting human gene transcription

**DOI:** 10.1101/290825

**Authors:** Naoki Osato

**Affiliations:** Department of Bioinformatic Engineering, Graduate School of Information Science and Technology, Osaka University, Osaka 565-0871, Japan

**Keywords:** transcriptional regulation, transcription factors, enhancer-promoter interactions, chromatin interactions, CTCF, open chromatin regions, histone modifications

## Abstract

**Background:** Chromatin interactions are essential in enhancer-promoter interactions (EPIs) and transcriptional regulation. CTCF and cohesin proteins located at chromatin interaction anchors and other DNA-binding proteins such as YY1, ZNF143, and SMARCA4 are involved in chromatin interactions. However, there is still no good overall understanding of proteins associated with chromatin interactions and insulator functions.

**Results:** Here, I describe a systematic and comprehensive approach for discovering DNA-binding motifs of transcription factors (TFs) that affect EPIs and gene expression. This analysis identified 96 biased orientations [64 forward-reverse (FR) and 52 reverse-forward (RF)] of motifs that significantly affected the expression level of putative transcriptional target genes in monocytes, T cells, HMEC, and NPC and included CTCF, cohesin (RAD21 and SMC3), YY1, and ZNF143; some TFs have more than one motif in databases; thus, the total number is smaller than the sum of FRs and RFs. KLF4, ERG, RFX, RFX2, HIF1, SP1, STAT3, and AP1 were associated with chromatin interactions. Many other TFs were also known to have chromatin-associated functions. The predicted biased orientations of motifs were compared with chromatin interaction data. Correlations in expression level of nearby genes separated by the motif sites were then examined among 53 tissues.

**Conclusion:** One hundred FR and RF orientations associated with chromatin interactions and functions were discovered. Most TFs showed weak directional biases at chromatin interaction anchors and were difficult to identify using enrichment analysis of motifs. These findings contribute to the understanding of chromatin-associated motifs involved in transcriptional regulation, chromatin interactions/regulation, and histone modifications.

## Background

Chromatin interactions play essential roles in enhancer-promoter interactions (EPIs) and transcriptional regulation. CTCF and cohesin proteins are located at the chromatin interaction anchors and facilitate the formation of loop structures. CTCF functions as an insulator by limiting the activity of enhancers in the loops (Fig. 1a) [1, 2]. The DNA-binding sequence of CTCF reveals its orientation bias at chromatin interaction anchors in terms of the forward-reverse (FR) orientation [1, 3, 4]. Other DNA-binding proteins such as ZNF143, YY1, and SMARCA4 (BRG1) are involved in chromatin interactions and EPIs [5–7]. ZNF143 is known to bind more often to promoters than distal regions, thereby establishing looping with the distal element bound by CTCF [5]. The DNA-binding motif of ZNF143 is enriched at chromatin interaction anchors (Z-score > 7), whereas the DNA-binding motif of YY1 is not enriched (Z-score < 2; [5] Figure 2a). CTCF, cohesin, ZNF143, YY1, and SMARCA4 are involved in various biological functions other than chromatin interaction. These transcription factors (TFs) bind to open chromatin regions near transcriptional start sites (TSS) as well, which may reduce the enrichment of the motifs at chromatin interaction anchors.

**Fig. 1.**
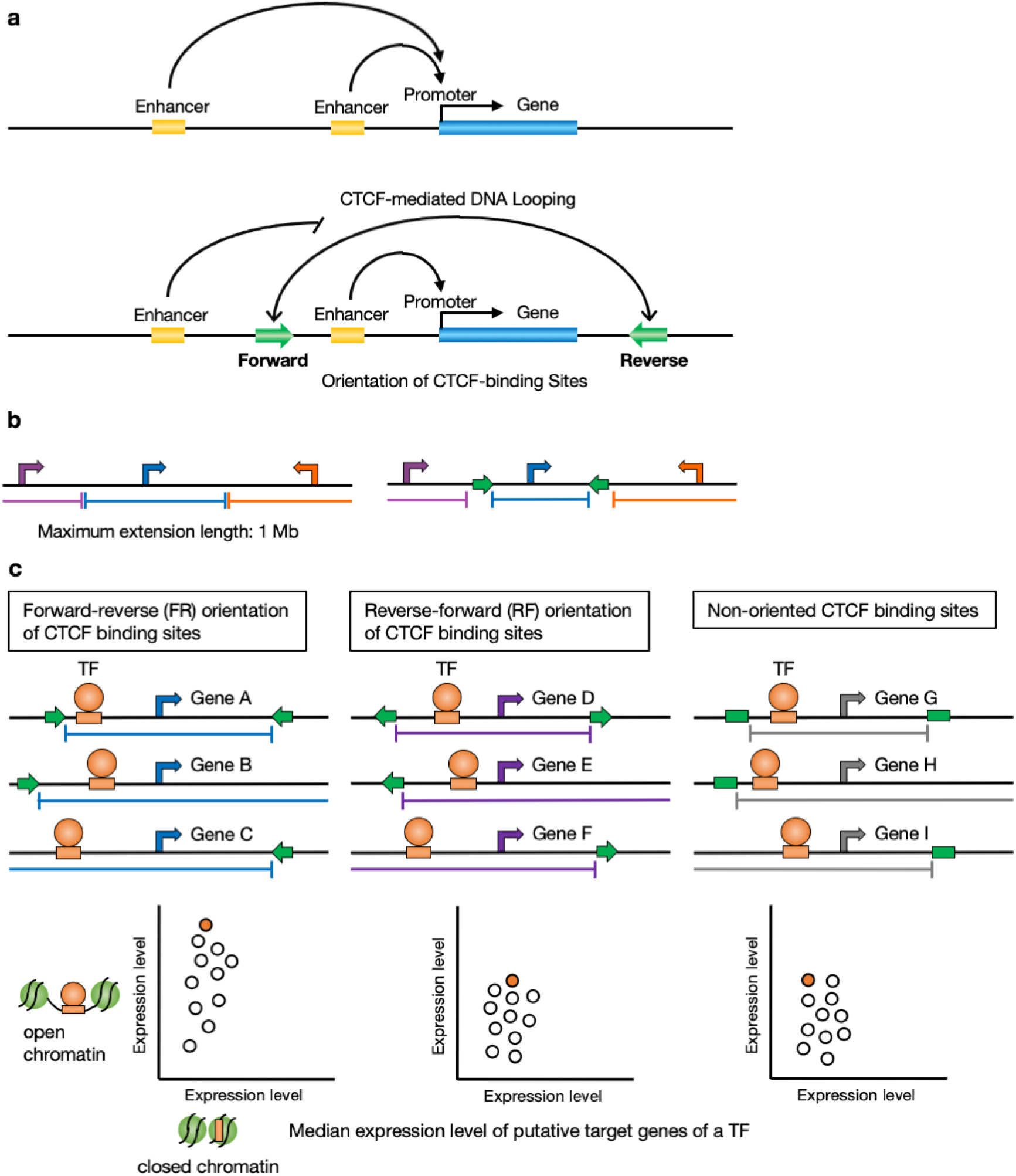
Biased orientations of DNA motifs of TFs affecting enhancer-promoter interactions. **a** Forward–reverse orientation of CTCF-binding sites is frequently found in chromatin interaction anchors. CTCF can block the interaction between enhancers and promoters limiting the activity of enhancers to certain functional domains [1–4]. **b** Computationally-defined regulatory domains for enhancer-promoter association (EPA) [12]. The *single nearest gene* association rule extends the regulatory domain to the midpoint between this gene’s TSS and the nearest gene’s TSS both upstream and downstream. The EPA domain was limited within the forward-reverse orientation of TFBSs of a TF in this study (e.g. CTCF in the figure). **c** Transcriptional target genes of each general TF bound in (i) distal open chromatin regions, which include enhancers, and (ii) closed chromatin regions in promoters were predicted using an EPA domain. An EPA domain was limited according to forward-reverse (FR), reverse-forward (RF) orientations, and non-oriented TFBSs of a putative insulator TF. Each dot in scatter plots represents the median expression level (FPKM) of target genes of a TF. The ratio of median expression level between (i) (y-axis) and (ii) (x-axis) was calculated to compare the distribution of the ratios of all general TFs among the three types of EPA domains.

**Fig. 2.**
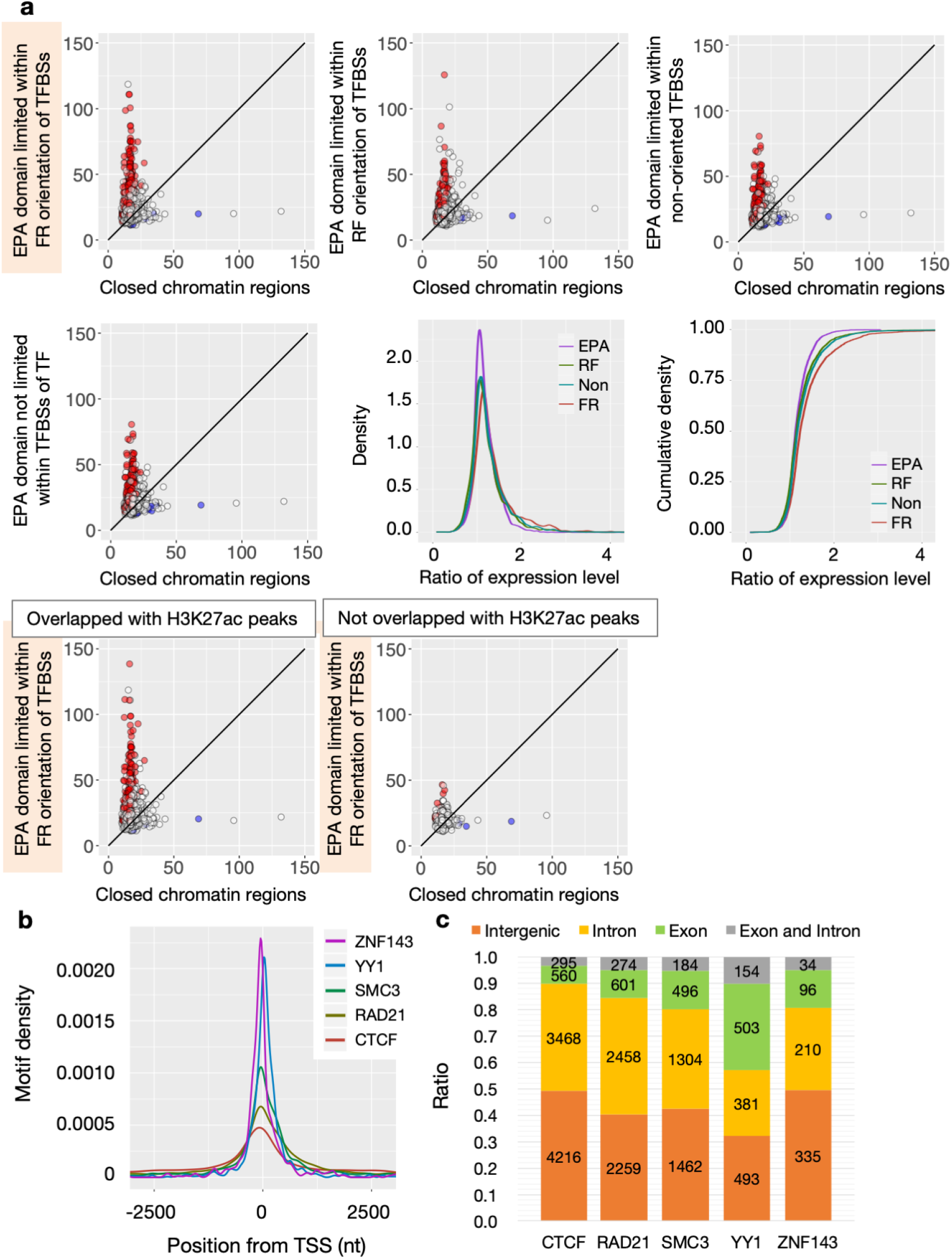
Identification of TFBSs in open chromatin regions and properties of TFBSs. **a** Comparison of expression level of putative transcriptional target genes. Each dot represents the median expression level (FPKM) of target genes predicted based on TFBSs of a TF in closed chromatin regions in promoters (x-axis) and an EPA domain limited within the FR orientation of CTCF-binding sites (y-axis) in T cells (upper left graph). The distribution of expression level was changed according to four criteria of EPA domains (four graphs) [11]. Red dots indicate the median expression level of target genes is significantly higher in an EPA domain than that in closed chromatin regions, and blue dots indicate the median expression level of target genes is significantly lower in an EPA domain than that in closed chromatin regions (Mann-Whitney test). The expression level of target genes predicted using an EPA domain tend to be higher than those predicted using closed chromatin regions, thereby implying that TFs serve as activators of target genes. The ratio of median expression level of target genes between an EPA domain and closed chromatin regions according to the four types of EPA domains is shown in terms of probability density and cumulative distribution (middle right two graphs). TFBSs overlapping H3K27ac ChIP-seq peaks were associated with a significant difference of expression level of target genes, and TFBSs not overlapping H3K27ac peaks were not associated with the difference (lower two graphs). This suggests that the significant difference of expression level is associated with the enhancer (and promoter) signals of H3K27ac. **b** Distribution of TFBSs of TFs near TSS. ZNF143 and YY1 tended to be located near TSS, and CTCF tended to be observed more in distant regions than those around TSS in monocytes. **c** Distribution of TFBSs of TFs in the human genome in monocytes. Most of TFBSs were found in intergenic and intron regions. YY1 is known to be at the TSS in the edge of exons of genes and is involved in transcriptional activation, repression, and initiation.

Computational methods to predict chromatin interactions have been developed [8, 9]. However, they were not intended to find DNA motifs of TFs involved in chromatin interactions. Xie et al. [10] proposed that the next challenge will be to develop systematic methods to discern the specific functions of these motifs in a genome-wide fashion.

DNA-binding proteins involved in chromatin interactions are considered to affect the transcription of genes within the loops formed by chromatin interactions. To confirm this hypothesis, I previously found that the expression level of human putative transcriptional target genes of a TF was significantly changed according to enhancer-promoter association (EPA) rules [11]. A regulatory domain for EPA was limited within the FR orientation of CTCF-binding sites, and the transcriptional target genes of each TF bound to enhancers and promoters were predicted using the EPA domain (Fig. 1b and c) [12]. The expression levels predicted using the EPA domain differed significantly from the expression levels predicted using promoters alone (or closed chromatin regions, where enhancers would be silenced, and nearby genes would express at low level [13]). The expression levels tended to increase in monocytes and CD4^+^ T cells, implying that TFs serve as activators of target genes. The expression levels tended to decrease in embryonic stem (ES) and induced pluripotent stem (iPS) cells, implying that TFs serve as repressors of target genes. The expression level of putative transcriptional target genes of a TF that binds enhancers is changed significantly when EPIs are predicted properly using the insulator sites of CTCF in different cell types. This observation suggests that the difference in the expression level reflects the accuracy of prediction of transcriptional target genes using a given EPA domain. Using this approach, I planned to search for other DNA-binding proteins involved in chromatin interactions, which affect the expression level of putative transcriptional target genes through their insulator function.

Experimental analyses of chromatin interactions in molecular biology have technical limitations. Chromatin interaction data are altered according to experimental techniques, the number of DNA sequencing reads, and even replication sets of the same cell type. Additionally, chromatin interaction data may not be abundant enough to contain all chromatin interactions and may include experimental noise such as ligation errors. To compensate the technical limitations of experimental analyses, a computational search was conducted for DNA motifs of TFs that affect EPIs and the expression level of putative transcriptional target genes in monocytes, T cells, human mammary epithelial cells (HMEC), and neural progenitor cells (NPC) without using chromatin interaction data. Then, I compared putative EPIs and transcription factor binding sites (TFBSs) with chromatin interaction and gene expression data.

## Results

### Search for biased orientations of DNA motifs

TFBSs were predicted using open chromatin regions and DNA motifs of TFs collected from various databases and journals (see Methods). Transcriptional target genes were predicted based on TFBSs in promoters and enhancers. An EPA domain was limited within the TFBSs with insulator function, such as CTCF and cohesin (RAD21 and SMC3). To find motifs of TFs with insulator function other than CTCF and cohesin, an EPA domain was limited within the TFBSs of a putative insulator TF (Fig. 1c). Transcriptional target genes of each general TF bound in (i) distal open chromatin regions, which include enhancers, and (ii) closed chromatin regions in promoters were predicted using the EPA domain in order to estimate the enhancer activity of each general TF. The ratio of median expression level of target genes of each general TF between (i) and (ii) was calculated. The distribution of the ratios of all general TFs was compared among forward-reverse (FR), reverse-forward (RF) orientations, and non-oriented (i.e., without considering orientation) groups of TFBSs using a two-sided Mann-Whitney test (*p*-value < 10^−7^). Putative insulator TFs with significant differences in the distributions of the ratios of expression level among FR, RF, and non-oriented were enumerated (Fig. 2a). TFBSs overlapping H3K27ac ChIP-seq peaks showed a significant difference in the expression level of their putative target genes between (i) and (ii), suggesting that the TFBSs were associated with enhancer (and promoter) signals of H3K27ac (Fig. 2a).

To confirm the predictions of TFBSs, the distributions of TFBSs were examined in the human genome. TFBSs involved in chromatin interactions were identified near TSS (Fig. 2b). ZNF143 is known to bind to the promoter of genes, thereby establishing looping with the distal element bound by CTCF [5]. YY1 is a multi-functional transcriptional regulator and is involved in transcriptional activation, repression, and initiation [14, 15]. The distributions of TFBSs seemed to be consistent with the features of the TFs. Most TFBSs were identified in intergenic and intron regions of the human genome (Fig. 2c).

Several hundreds of FR and RF orientations were found in monocytes, T cells, HMEC, and NPC, which included DNA motifs with weak biased orientations (Fig. 3). The number of TFs was likely to be changed, according to the number of DNase-seq reads, since the number of the reads in monocytes was larger than those of the other three cell types. When DNase-seq reads increase, more TFBSs may be predicted to some extent. Notably, 96 biased (64 FR and 52 RF) orientations of DNA-binding motifs of TFs were discovered and found to be common between the four cell types, whereas non-oriented DNA-binding motifs of TFs were not detected (Fig. 3; Table 1; Additional file 1: Table S1). The FR orientation of DNA motifs included CTCF, cohesin (RAD21 and SMC3), YY1, and ZNF143, which are involved in chromatin interactions and EPIs. Some TFs have both FR and RF orientations of DNA motifs; thus, the total number 96 is smaller than the sum of FRs and RFs. Palindromic DNA motifs of TFs do not have FR and RF distinction; thus, they do not show a directional bias.

**Fig. 3.**
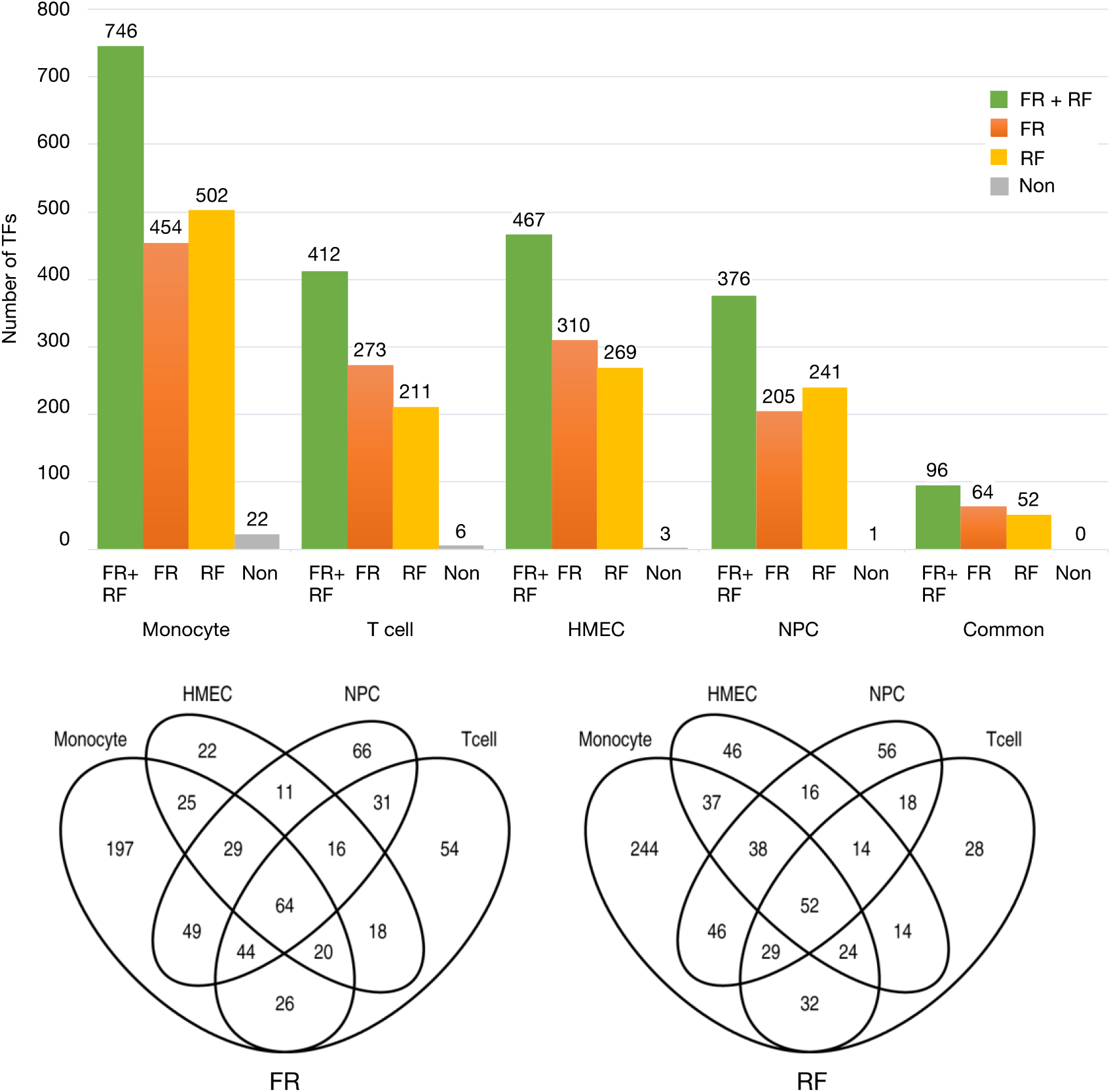
Biased orientations of TFBSs of TFs affect the transcription of genes by the putative insulator function of the TFs. Ninety-six (sixty-four FR and fifty-two RF) biased orientations of TFBSs of TFs showed significant differences in the distributions of the ratios of expression level among FR, RF, and non-orientation in monocytes, T cells, HMEC, and NPC, whereas non-oriented TFBSs of TFs did not show significant differences in expression level.

**Table 1.**
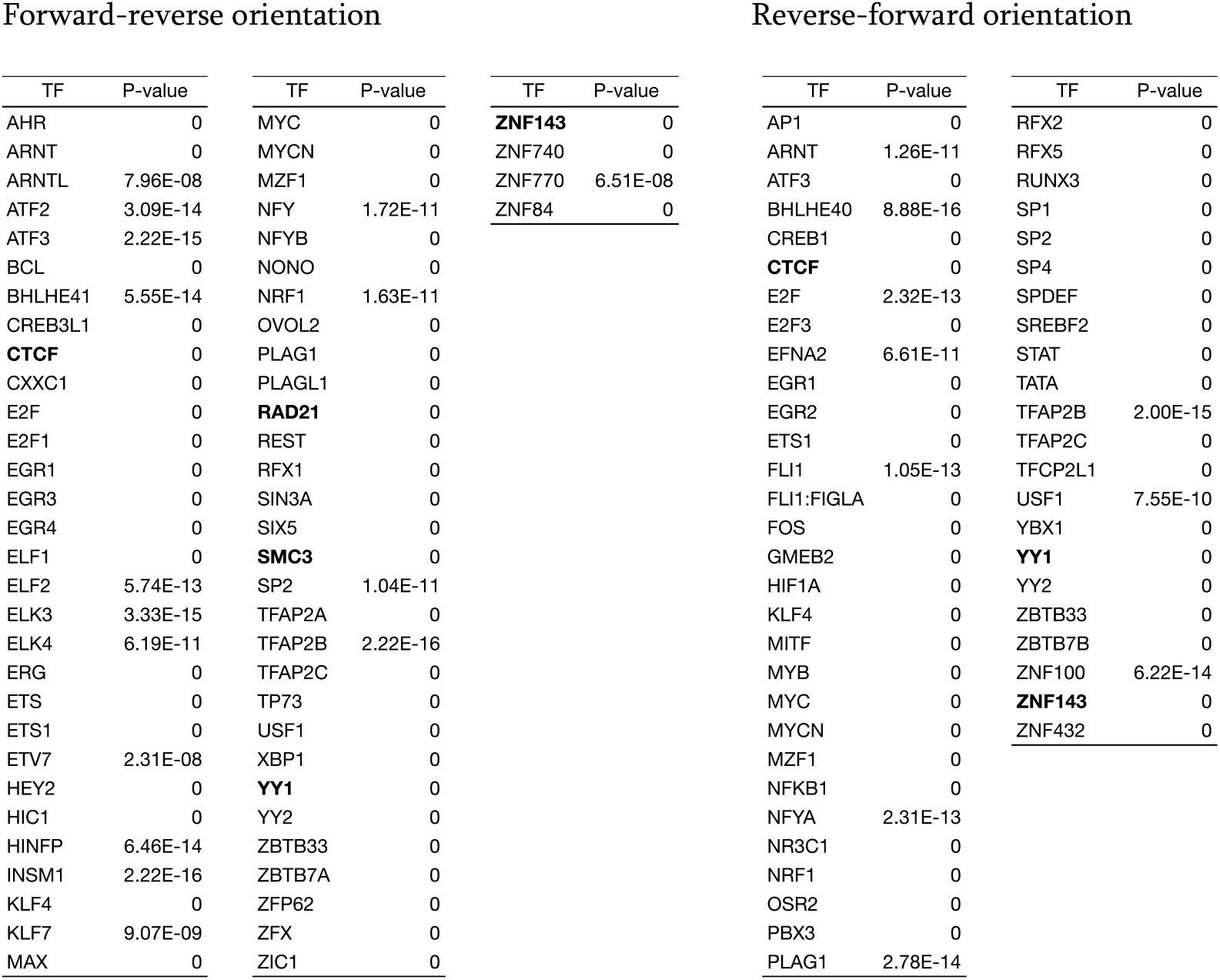
Biased orientations of DNA-binding motifs of TFs in common between monocytes, T cells, HMEC, and NPC. TF: DNA-binding motifs of transcription factors.

Among the 96 TFs, KLF4 [16], ERG [17, 18], RFX [19], RFX2 [20], HIF1 [21], SP1 [22], STAT3 [23], and AP1 [24] were found to be associated with chromatin interactions. An important example is KLF4, which is associated with three-dimensional enhancer rewiring and transcriptional changes during the reprogramming of mouse embryonic fibroblasts to pluripotent stem cells [16]. ATF3 [25], ZBTB7A [26], NRF1 [6, 27], TFAP2 [28], Myc [29, 30], the ETS family, ELK3 [31], ELK4 [32], RUNX3 [33], USF1 [34, 35], AHR/ARNT [36], SP2 [37], EGR2 [38–40], ZBTB33 [41], NF-Y [42, 43], MITF [44, 45], NONO [46], FLI1 [47], E2F [48–50], BCL [51], CXXC1 [52], E2F1 [53], EGR1 [54], REST [55, 56], RFX [57], SIN3A [58–60], AP1 [61], and STAT3 and STAT5 [62] were associated with chromatin remodeling, chromatin accessibility, pioneer factor, and/or histone modification. TFBSs of the TFs other than those involved in chromatin interactions would affect gene expression by chromatin-associated functions. Huang et al. [34] have mentioned that the insulator element at the 5’ end of the chicken beta-globin locus acts as a barrier, thus protecting transgenes against silencing effects of adjacent heterochromatin. The transcription factor USF1 binds within the insulator at a site important in adjacent nucleosomes for generating histone modifications associated with active chromatin and, by inference, with barrier function.

The biased orientations of TFBSs might be located near those of known TFs involved in chromatin interactions such as CTCF, RAD21, SMC3, YY1, and ZNF143; thus, they showed significant differences in the expression level of putative transcriptional target genes among FR, RF, and non-orientation. However, among 447 FR and 496 RF orientations of DNA motifs in monocytes, the TFBSs of only 16 (4%) FR and 28 (6%) RF orientations were located within 1 kb of ≥ 20% of the TFBSs of either of the known TFs and their associated TFs such as CTCFL and YY2 (Additional file 1: Table S2).

To examine whether the biased orientations of TFBSs were associated with CTCF-binding sites, an EPA domain was limited within the TFBSs of biased orientations of a TF in the 96 TFs found to be common among the four cell types. Then, transcriptional target genes of each general TF were predicted from (i) open chromatin regions in the EPA domain that did not include CTCF-binding sites upstream and downstream of a gene, and (ii) closed chromatin regions in promoters. The ratio of expression level between (i) and (ii) was calculated. The biased orientations of CTCF consist of 19 DNA-binding motifs in monocytes. When upstream and downstream of genes with the TFBSs of each DNA-binding motif sequence of CTCF were eliminated, 93, 92, 93, and 84 TFs out of 95, except for CTCF, showed a significant difference in the distribution of the ratios of expression level (*p*-value < 10^−7^) in monocytes, CD4^+^ T cells, HMEC, and NPC, respectively (Additional file 1: Table S3). This suggests that the biased orientations of TFBSs are not significantly associated with CTCF binding-sites.

### Comparison with chromatin interaction data

To examine whether the biased orientations of DNA motifs were associated with chromatin interactions, EPIs predicted using an EPA domain were compared with chromatin interaction data, using three replications (B2T1, B2T2, and B3T1) of H3K27ac HiChIP chromatin interaction data in CD4^+^ T cells separately [63]. The resolutions of HiChIP chromatin interaction data and EPIs were adjusted to 5 kilobases (kb). The number of unique HiChIP cis (i.e., in the same chromatin) chromatin interactions was 19,926,360 at 5-kb resolution in B2T1 replication of T cells; 666,149 at 5-kb resolution with ≥1,000 counts for each interaction; and 78,209 at 5-kb resolution with ≥6,000 counts for each interaction (Additional file 1: Table S4). EPIs were predicted using three types of EPA domains: (i) EPA domain limited within the FR or RF orientation of TFBSs of a TF with putative insulator function such as CTCF, (ii) EPA domain limited within the TFBSs of a TF without considering their orientation, and (iii) EPA domain which was not limited within TFBSs. Despite the limited resolution of the chromatin interaction data, a total of 201 biased orientations [133 of 273 (49%) FR and 90 of 211 (43%) RF; some TFs have both FR and RF motifs; thus, the total number is smaller than the sum of FRs and RFs] of DNA motifs in T cells included CTCF, cohesin (RAD21 and SMC3), ZNF143, and YY1 in three replications (B2T1, B2T2, and B3T1). For the DNA motifs, a significantly higher ratio of EPIs overlapped with HiChIP chromatin interactions with ≥1,000 counts for each interaction in EPA domain (i) than in EPA domain (ii) and (iii) in T cells (Table 2). For a total of 390 biased orientations [261 of 273 (96%) FR and 199 of 211 (94%) RF] of DNA motifs in T cells, a significantly higher ratio of EPIs overlapped with the chromatin interactions in EPA domain (i) than EPA domain (iii) (Table 2; Additional file 1: Table S4 and S5). The difference between EPIs predicted using EPA domain (i) and (ii) seemed to be difficult to distinguish using the chromatin interaction data and a statistical test in some cases. However, for the difference between EPIs predicted using EPA domain (i) and (iii), a larger number of biased orientations of DNA motifs were correlated with chromatin interaction data. Most (95%) of the biased orientations of DNA motifs were correlated with chromatin interactions when comparing EPIs predicted using EPA domain (i) and (iii) with HiChIP chromatin interactions. Chromatin interaction data were obtained from different samples of DNase-seq open chromatin regions; thus, individual differences may exist. Individual differences of chromatin interactions were larger than those of open chromatin regions [63]. Additionally, the analyses for Th17, Treg, and GM12878 were shown in Additional file 2.

**Table 2.** Comparison of enhancer-promoter interactions (EPIs) with chromatin interactions in T cells. The upper tables show the number of biased orientations of DNA motifs, where a significantly higher ratio of EPIs that were predicted using an EPA domain (i) overlapped with HiChIP chromatin interactions than EPA domain (ii) and (ii) in T cells. The lower tables show that among 64 FR and 52 RF orientations of DNA motifs found in common in monocytes, T cells, HMEC, and NPC, 44 (69%) FR and 28 (54%) RF were matched with the analysis of HiChIP chromatin interactions according to three types of EPA domains. TF: DNA-binding motifs of a transcription factor. Score: −log_10_ (*p*-value). Score 1: Comparison of EPA domain (i) (an EPA domain limited within the FR or RF orientation of TFBSs) with EPA domain (iii) (an EPA domain not limited). Score 2: Comparison of EPA domain (i) with EPA domain (ii) (an EPA domain limited within the non-oriented TFBSs). FR and RF orientations of DNA motifs found to be common in the four cell types.

**1. Comparison of EPI with HiChIP data among three EPA (i), (ii), and (iii)**
| Replication of HiChIP data | Total no. (FR + RF)<br>of DNA motifs | No. of FR<br>DNA motifs | No. of RF<br>DNA motifs |
| --- | --- | --- | --- |
| B2T1 | 226 | 153 | 101 |
| B2T2 | 221 | 149 | 98 |
| B3T1 | 237 | 160 | 109 |
| DNA motifs found in common among replications |  |  |  |
| B2T1 and B2T2 | 212 | 141 | 95 |
| B2T1 and B3T1 | 212 | 143 | 93 |
| B2T2 and B3T1 | 208 | 138 | 93 |
| B2T1, B2T2, and B3T1 | 201 | 133 | 90 |

**2. Comparison of EPI with HiChIP data between two EPA (i) and (iii)**
| Replication of HiChIP data | Total no. (FR + RF)<br>of DNA motifs | No. of FR<br>DNA motifs | No. of RF<br>DNA motifs |
| --- | --- | --- | --- |
| B2T1 | 395 | 262 | 205 |
| B2T2 | 398 | 264 | 204 |
| B3T1 | 394 | 263 | 203 |
| DNA motifs found in common among replications |  |  |  |
| B2T1 and B2T2 | 394 | 262 | 202 |
| B2T1 and B3T1 | 391 | 261 | 202 |
| B2T2 and B3T1 | 392 | 262 | 200 |
| B2T1, B2T2, and B3T1 | 390 | 261 | 199 |

| TF | Score 1 | Score 2 | TF | Score 1 | Score 2 | TF | Score 1 | Score 2 |
| --- | --- | --- | --- | --- | --- | --- | --- | --- |
| AHR | 38.81 | 13.41 | **RAD21** | 258.21 | 4.81 | ARNT | 323.31 | 14.12 |
| ARNTL | 155.78 | 1.96 | RFX1 | 159.78 | 17.05 | ATF3 | 323.31 | 14.64 |
| ATF3 | 323.31 | 35.22 | **SMC3** | 222.09 | 1.76 | BHLHE40 | 323.31 | 12.42 |
| BCL | 153.72 | 3.52 | SP2 | 94.11 | 9.54 | **CTCF** | 240.61 | 4.56 |
| CREB3L1 | 323.31 | 27.96 | TFAP2A | 25.86 | 20.26 | E2F | 78.16 | 7.07 |
| **CTCF** | 323.31 | 3.50 | TFAP2B | 90.03 | 6.10 | E2F3 | 65.67 | 8.85 |
| CXXC1 | 69.92 | 3.30 | TFAP2C | 169.60 | 8.55 | EFNA2 | 255.01 | 10.95 |
| E2F | 122.12 | 21.87 | TP73 | 198.45 | 7.75 | GMEB2 | 323.31 | 7.81 |
| E2F1 | 43.22 | 2.85 | USF1 | 194.06 | 3.56 | KLF4 | 213.35 | 9.49 |
| EGR1 | 79.09 | 3.79 | XBP1 | 323.31 | 44.58 | MYB | 323.31 | 17.68 |
| EGR3 | 85.25 | 34.18 | **YY1** | 62.59 | 5.30 | MYC | 323.31 | 12.23 |
| ELF1 | 203.38 | 3.86 | ZBTB7A | 52.89 | 30.58 | MZF1 | 200.49 | 22.73 |
| ELF2 | 227.62 | 7.83 | ZFX | 13.33 | 14.55 | NFYA | 323.31 | 11.75 |
| ELK4 | 117.83 | 4.81 | **ZNF143** | 323.31 | 35.38 | NR3C1 | 43.16 | 8.31 |
| ERG | 199.81 | 25.70 |  |  |  | OSR2 | 111.77 | 14.65 |
| ETS | 196.80 | 1.63 |  |  |  | SP2 | 294.86 | 2.59 |
| ETS1 | 308.00 | 3.92 |  |  |  | SP4 | 110.94 | 19.63 |
| HIC1 | 223.55 | 4.56 |  |  |  | SPDEF | 185.85 | 14.35 |
| INSM1 | 193.85 | 15.87 |  |  |  | SREBF2 | 297.27 | 1.62 |
| KLF4 | 209.71 | 9.38 |  |  |  | TATA | 2.16 | 4.02 |
| KLF7 | 229.95 | 11.96 |  |  |  | TFAP2B | 29.46 | 12.62 |
| MAX | 323.31 | 8.35 |  |  |  | TFAP2C | 112.10 | 51.21 |
| MYC | 84.79 | 24.93 |  |  |  | **YY1** | 323.31 | 9.92 |
| MYCN | 165.16 | 25.87 |  |  |  | YY2 | 47.49 | 2.61 |
| MZF1 | 81.83 | 2.29 |  |  |  | ZBTB33 | 90.29 | 7.78 |
| NFY | 323.31 | 3.42 |  |  |  | ZBTB7B | 43.90 | 23.59 |
| NONO | 323.31 | 9.24 |  |  |  | ZNF100 | 27.67 | 4.75 |
| NRF1 | 45.13 | 30.75 |  |  |  | **ZNF143** | 63.03 | 9.69 |
| OVOL2 | 323.31 | 31.61 |  |  |  |  |  |  |
| PLAG1 | 177.90 | 11.02 |  |  |  |  |  |  |

In situ Hi-C data in HMEC, which are available in a 4D nucleome data portal, were compared with putative EPIs. The number of unique in situ Hi-C cis chromatin interactions was 121,873,642 at 3-kb resolution. For a total of 222 biased orientations [155 of 310 (50%) FR and 108 of 269 (40%) RF] of DNA motifs, which included CTCF, cohesin (RAD21 and SMC3), ZNF143, and YY1, a significantly higher ratio of EPIs overlapped with in situ Hi-C chromatin interactions (≥1 count for each interaction, i.e., all interactions) in EPA domain (i) than in EPA domain (ii) and (iii) (Additional file 1: Table S7). When comparing EPIs predicted using only EPA domain (i) and (iii) with the chromatin interactions, for a total of 406 biased orientations [268 of 310 (86%) FR and 230 of 269 (86%) RF] of DNA motifs, a significantly higher ratio of EPIs overlapped with the chromatin interactions in EPA domain (i) than EPA domain (iii) (Additional file 1: Table S7). There were six replications of in situ Hi-C experimental data of HMEC, but other replications contained much smaller numbers of chromatin interactions. The HiChIP chromatin interaction data in T cells were expected to be enriched with EPIs, since they were produced with H3K27ac antibody to capture chromatin interactions. In situ Hi-C data consist of all types of chromatin interactions and thus contain a smaller number of EPIs than HiChIP data. Chromatin interactions with a small number of counts may be artifacts caused by ligation errors during experiments [64, 65]. Therefore, other replications of in situ Hi-C data with a smaller number of chromatin interactions will not be available for this analysis.

Promoter capture Hi-C data of NPC were downloaded and analyzed by the same procedure as that for T cells and HMEC [66]. The number of unique promoter capture Hi-C cis chromatin interactions was 9,436,689 at 5-kb resolution. For a total of 109 biased orientations [61 of 205 (28%) FR and 60 of 241 (25%) RF] of DNA motifs, which included CTCF, cohesin (RAD21 and SMC3), ZNF143, and YY1, a significantly higher ratio of EPIs overlapped with promoter capture Hi-C chromatin interactions (≥1 count for each interaction, i.e. all interactions) in EPA domain (i) than in EPA domain (ii) and (iii) (Additional file 1: Table S8). When comparing EPIs predicted using only EPA domain (i) and (iii) with the chromatin interactions, for a total of 186 biased orientations [99 of 205 (48%) FR and 103 of 241 (43%) RF] of DNA motifs, a significantly higher ratio of EPIs overlapped with the chromatin interactions in EPA domain (i) than EPA domain (iii) (Additional file 1: Table S8).

All three types of chromatin interaction data (HiChIP, in situ Hi-C, and promoter capture Hi-C) indicated that EPIs predicted using EPA domain (i) that was limited within TFBSs of known TFs involved in chromatin interactions (CTCF, RAD21, SMC3, ZNF143 and YY1) overlapped with more chromatin interactions than those using EPA domain (ii) and (iii) in six cell types (T cells, Th17, Treg, GM12878, HMEC, and NPC) (the analyses for Th17, Treg, and GM12878 were shown in Additional file 2). In monocytes, Hi-C data was available, but it was not analyzed due to the low resolution of chromatin interaction data (50 kb).

The biased orientations of TFBSs of known TFs involved in chromatin interactions such as CTCF, RAD21, SMC3, YY1, and ZNF143 might overlap with the biased orientations of TFBSs of other TFs; thus, they showed a significantly higher ratio of EPIs overlapped with chromatin interactions in EPA domain (i) than EPA domain (ii) and (iii). However, among 273 FR and 211 RF orientations of DNA motifs of TFs not including the known TFs in T cells, the TFBSs of only 15 (6%) FR and 9 (4%) RF orientations of DNA motifs of TFs overlapped with ≥ 60% of the TFBSs of biased orientations of DNA motifs of either of the known TFs at 5-kb resolution.

Moreover, to investigate the enhancer activity of EPIs, the distribution of expression level of putative target genes of EPIs was compared between EPIs that overlapped with HiChIP chromatin interactions and EPIs that did not overlap with them. Though the target genes of EPIs were selected from top 4,000 transcripts (genes) in terms of the expression level of all transcripts (genes) excluding transcripts not expressed in T, Th17, Treg, and GM12878 cells separately, target genes of EPIs that overlapped with chromatin interactions showed a significantly higher expression level than non-overlapped EPIs, suggesting that EPIs that overlapped with chromatin interactions activated the expression of target genes in the four cell types. For almost all (99%-100%) FR and RF orientations of DNA motifs, a significantly higher expression level of putative target genes was observed for EPIs that overlapped with chromatin interactions than for EPIs that did not overlap in the four cell types (Additional file 1: Table S5 and S6). DNA motifs showing a significantly lower expression level were not observed in this analysis. HiChIP data were produced using H3K27ac antibody; thus, chromatin interactions acting as repressors would not be identified. However, for in situ Hi-C and promoter capture Hi-C data in HMEC and NPC, the significant difference in the expression level of putative target genes of EPIs was observed in a small percentage (less than 1%) of biased orientations of DNA motifs.

Recently, high resolution chromatin interaction data became publicly available [67]. The biased orientations of TFBSs of TFs were examined using Micro-C chromatin interaction data in human fibroblasts. The resolution of the data was adjusted to 200 bp. TFBSs of CTCF, RAD21, SMC3, and SMRC2 showed a significant directional bias at the pairs of chromatin interaction anchors (*p*-value < 10^−50^). SMARCA4 is known to be involved in higher-order chromatin structures such as the topologically associating domain (TAD) [7]. DNA motifs of SMARCA4 did not exist in the databases of DNA motifs of TFs used in this analysis. SMRC2 was not identified in this analysis because it may be involved in higher-order chromatin structures that do not affect EPI and gene expression directly by its insulator function. ATF3 and ZBTB7A in Table 1 were found among the top 32 biased orientations of TFBSs of TFs at the pairs of chromatin interaction anchors with *p*-value < 10^−50^ (Additional file 1: Table S9). Deletion of an ATF3-binding site affected the expression of a target gene, which was far apart from the site [25]. ZBTB7A is involved in the changes in chromatin accessibility and the regulation of gene expression [26]. TFBSs of YY1 and ZNF143 also indicated a weak directional bias at the chromatin interaction anchors (Additional file 1: Table S9). In this analysis, biased orientations of TFBSs of TFs were investigated based on the difference of distribution of the expression level of putative transcriptional target genes. YY1 and ZNF143 were found to have significant directional biases, thereby suggesting that the analysis in this study could find TFs that did not have a strong directional bias based on the number of TFBSs at chromatin interaction anchors. The other TFs, except for CTCF, RAD21, SMC3, ATF3, and ZBTB7A in Table 1, showed weak directional biases at the chromatin interaction anchors (Additional file 1: Table S9).

### Correlation of expression level of gene pairs separated by TFBS

To investigate whether the biased orientations of TFBSs affected gene expression, the expression level of protein-coding genes, that are most closely located upstream and downstream of any biased orientations of TFBSs in the human genome, was compared in monocytes. The expression levels of closely located genes including divergent gene pairs tend to be correlated among tissues [68, 69]. When a biased orientation of TFBSs serves as an insulator (e.g. CTCF), the correlation of gene expression level would be reduced. Among 96 biased (64 FR and 52 RF) orientations of TFBSs found in common in monocytes, T cells, HMEC, and NPC, TFs with ≥50 genomic locations of the TFBSs were selected after the elimination of ubiquitously expressed genes among 53 tissues and TFBSs near the genes with a coefficient of variance (CV) < 90. After this filtering, 43 of 60 (72%) FR and 40 of 50 (80%) RF orientations showed significantly lower correlations of expression levels of gene pairs separated by the TFBSs of a TF than the correlations of all pairs of neighboring genes with intergene distance <1 megabase (Mb) (Mann-Whitney test, *p*-value < 0.05) (Table 3). The FR and RF orientations of TFs included known TFs involved in chromatin interactions such as CTCF, RAD21, SMC3, YY1, and ZNF143. The top 20 FR and RF orientations of DNA motifs were selected in ascending order of the median correlation coefficient of expression levels of gene pairs separated by the TFBSs of a TF. The 20 TFs included known TFs involved in chromatin interactions such as CTCF, RAD21, and SMC3 (Fig. 4) (Mann-Whitney test, *p*-value < 0.01). Instead of using all TFBSs of a TF, when TFBSs of a TF were limited to sites used in the analysis of biased orientations of DNA motifs in an EPA domain, 33 of 55 (60%) FR and 36 of 47 (77%) RF orientations showed significantly lower correlations of expression levels of gene pairs separated by the TFBSs (Additional file 1: Table S10). With intergene distance < 20 kb, 21 of 62 (34%) FR and 8 of 52 (15%) RF orientations showed significantly lower correlations of expression levels of gene pairs separated by the TFBSs of a TF than the correlations of all pairs of neighboring genes (Additional file 1: Table S10).

**Table 3.**
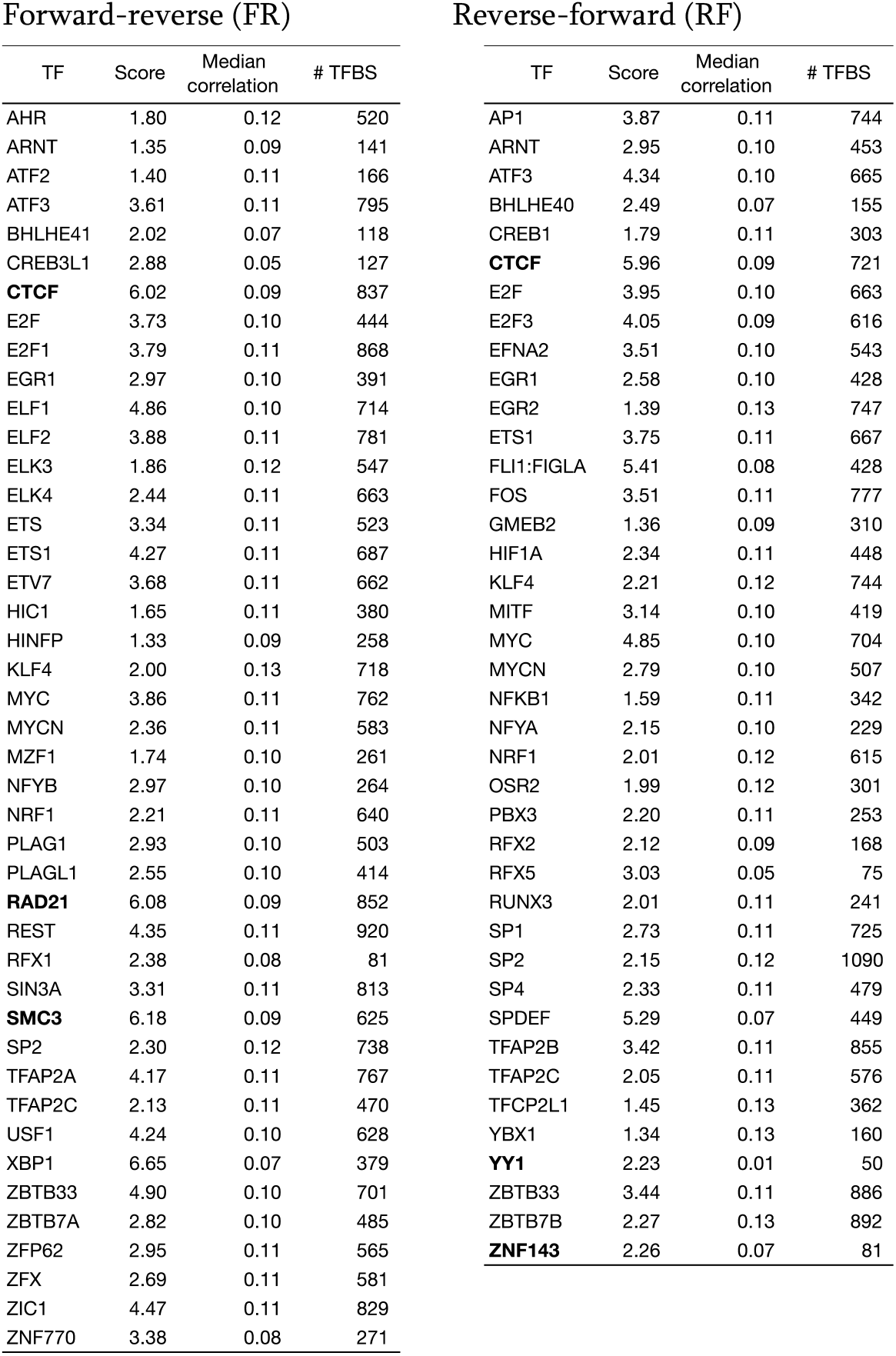
Correlation of the expression level of gene pairs separated by TFBSs of a TF. Pairs of genes separated by 64 FR and 52 RF orientations of TFBSs of TFs found to be common between monocytes, T cells, HMEC, and NPC were analyzed to find significantly lower correlation of their expression level compared to the correlation of all pairs of nearby genes among 53 tissues. Score: −log_10_ (*p*-value) of the Mann-Whitney test. Median correlation: median correlation coefficient of expression level of nearby gene pairs separated by TFBSs. The median correlation coefficient of expression level of all pairs of nearby genes was 0.17. # TFBS: number of TFBSs of a TF separating gene pairs with expression data. All TFBSs of a TF were used, except for TFBSs separating pairs of ubiquitously expressed genes with coefficient of variance < 90.

| Forward-reverse (FR) |  |  |  | Reverse-forward (RF) |  |  |  |
| --- | --- | --- | --- | --- | --- | --- | --- |
| TF | Score | Median correlation | # TFBS | TF | Score | Median correlation | # TFBS |
| AHR | 1.80 | 0.12 | 520 | AP1 | 3.87 | 0.11 | 744 |
| ARNT | 1.35 | 0.09 | 141 | ARNT | 2.95 | 0.10 | 453 |
| ATF2 | 1.40 | 0.11 | 166 | ATF3 | 4.34 | 0.10 | 665 |
| ATF3 | 3.61 | 0.11 | 795 | BHLHE40 | 2.49 | 0.07 | 155 |
| BHLHE41 | 2.02 | 0.07 | 118 | CREB1 | 1.79 | 0.11 | 303 |
| CREB3L1 | 2.88 | 0.05 | 127 | **CTCF** | 5.96 | 0.09 | 721 |
| **CTCF** | 6.02 | 0.09 | 837 | E2F | 3.95 | 0.10 | 663 |
| E2F | 3.73 | 0.10 | 444 | E2F3 | 4.05 | 0.09 | 616 |
| E2F1 | 3.79 | 0.11 | 868 | EFNA2 | 3.51 | 0.10 | 543 |
| EGR1 | 2.97 | 0.10 | 391 | EGR1 | 2.58 | 0.10 | 428 |
| ELF1 | 4.86 | 0.10 | 714 | EGR2 | 1.39 | 0.13 | 747 |
| ELF2 | 3.88 | 0.11 | 781 | ETS1 | 3.75 | 0.11 | 667 |
| ELK3 | 1.86 | 0.12 | 547 | FLI1:FIGLA | 5.41 | 0.08 | 428 |
| ELK4 | 2.44 | 0.11 | 663 | FOS | 3.51 | 0.11 | 777 |
| ETS | 3.34 | 0.11 | 523 | GMEB2 | 1.36 | 0.09 | 310 |
| ETS1 | 4.27 | 0.11 | 687 | HIF1A | 2.34 | 0.11 | 448 |
| ETV7 | 3.68 | 0.11 | 662 | KLF4 | 2.21 | 0.12 | 744 |
| HIC1 | 1.65 | 0.11 | 380 | MITF | 3.14 | 0.10 | 419 |
| HINFP | 1.33 | 0.09 | 258 | MYC | 4.85 | 0.10 | 704 |
| KLF4 | 2.00 | 0.13 | 718 | MYCN | 2.79 | 0.10 | 507 |
| MYC | 3.86 | 0.11 | 762 | NFKB1 | 1.59 | 0.11 | 342 |
| MYCN | 2.36 | 0.11 | 583 | NFYA | 2.15 | 0.10 | 229 |
| MZF1 | 1.74 | 0.10 | 261 | NRF1 | 2.01 | 0.12 | 615 |
| NFYB | 2.97 | 0.10 | 264 | OSR2 | 1.99 | 0.12 | 301 |
| NRF1 | 2.21 | 0.11 | 640 | PBX3 | 2.20 | 0.11 | 253 |
| PLAG1 | 2.93 | 0.10 | 503 | RFX2 | 2.12 | 0.09 | 168 |
| PLAGL1 | 2.55 | 0.10 | 414 | RFX5 | 3.03 | 0.05 | 75 |
| **RAD21** | 6.08 | 0.09 | 852 | RUNX3 | 2.01 | 0.11 | 241 |
| REST | 4.35 | 0.11 | 920 | SP1 | 2.73 | 0.11 | 725 |
| RFX1 | 2.38 | 0.08 | 81 | SP2 | 2.15 | 0.12 | 1090 |
| SIN3A | 3.31 | 0.11 | 813 | SP4 | 2.33 | 0.11 | 479 |
| **SMC3** | 6.18 | 0.09 | 625 | SPDEF | 5.29 | 0.07 | 449 |
| SP2 | 2.30 | 0.12 | 738 | TFAP2B | 3.42 | 0.11 | 855 |
| TFAP2A | 4.17 | 0.11 | 767 | TFAP2C | 2.05 | 0.11 | 576 |
| TFAP2C | 2.13 | 0.11 | 470 | TFCP2L1 | 1.45 | 0.13 | 362 |
| USF1 | 4.24 | 0.10 | 628 | YBX1 | 1.34 | 0.13 | 160 |
| XBP1 | 6.65 | 0.07 | 379 | **YY1** | 2.23 | 0.01 | 50 |
| ZBTB33 | 4.90 | 0.10 | 701 | ZBTB33 | 3.44 | 0.11 | 886 |
| ZBTB7A | 2.82 | 0.10 | 485 | ZBTB7B | 2.27 | 0.13 | 892 |
| ZFP62 | 2.95 | 0.11 | 565 | **ZNF143** | 2.26 | 0.07 | 81 |
| ZFX | 2.69 | 0.11 | 581 |  |  |  |  |
| ZIC1 | 4.47 | 0.11 | 829 |  |  |  |  |
| ZNF770 | 3.38 | 0.08 | 271 |  |  |  |  |

**Fig. 4.**
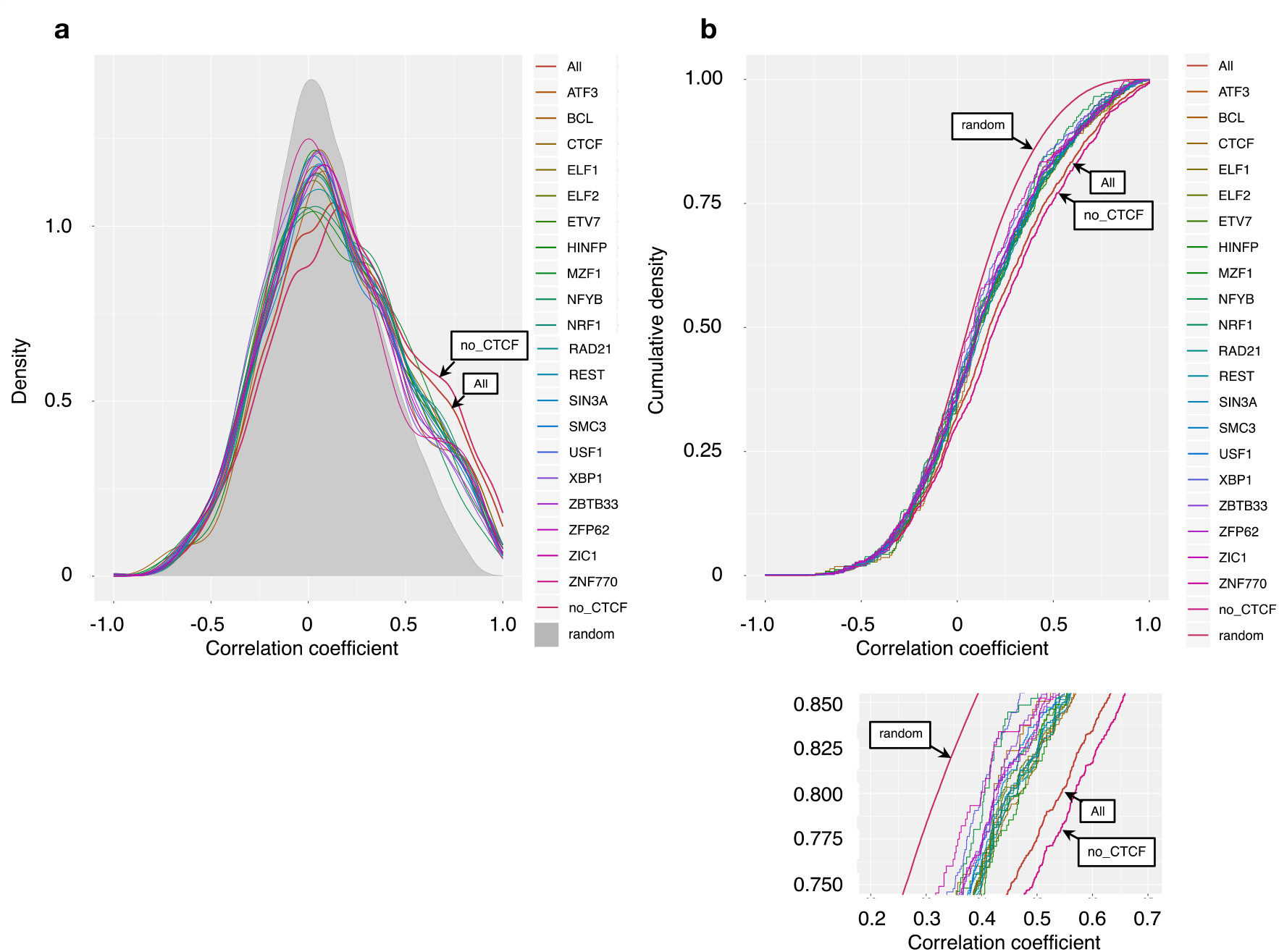
Genes separated by predicted TFBSs of biased orientations of TFs are less correlated in gene expression. The top 20 TFs are selected according to ascending order of the median correlation coefficient. The correlation coefficient between neighboring gene pairs is shown in terms of probability density **a** and cumulative distribution **b**. The brown line (All) represents the correlation between all neighboring genes; gray shading in **a** and red line (random) in **b** represents correlation between randomly chosen gene pairs; and the purple line (no_CTCF) represents the correlation between gene pairs not separated by predicted CTCF-binding sites.

A large number of TFBSs of a TF tend to show significantly lower correlations of gene expression (median numbers of TFBSs of a TF showing significantly lower correlations were 656 and 479 for FR and RF, respectively, and median numbers of TFBSs of a TF not showing significantly lower correlations were 122 and 115 for FR and RF, respectively. The Mann-Whitney test *p*-value = 0 and < 10^−10^ for FR and RF, respectively. This suggests that if the criteria to examine the correlations of expression levels of gene pairs separated by TFBSs are more stringent, and the number of TFBSs is reduced, it would be more difficult to use a statistical test to compare the correlations of expression levels of gene pairs separated by the reduced TFBSs, particularly for TFs with a small number of TFBSs.

Among 91 biased orientations of DNA motifs of TFs found in common among the four cell types, after excluding the five known TFs involved in chromatin interactions such as CTCF, RAD21, SMC3, YY1, and ZNF143, TFBSs of 88 TFs were located at ≥1 genomic regions between gene pairs not including TFBSs of the known TFs and at ≥1 genomic regions between low correlation (< 0.13) of expression level of gene pairs (Table 3). TFBSs of 52 TFs were placed at ≥10 genomic regions, though some proportion of TFBSs of the known TFs might not be associated with insulator function.

### Co-location of motifs with biased orientations

To examine the association of directional DNA-binding motifs, co-location of motifs within 200 bp (approximately the minimum length of an open chromatin region of 146 bp) of each other was analyzed in monocytes, T cells, HMEC, and NPC. The same pairs of motifs upstream and downstream of genes in EPA domain were enumerated, and the pairs of motifs found in ≥100 genomic regions were listed (Table 4; Additional file 1: Table S11). A majority of the pairs of motifs overlapped with more than 1 base, but the same tendency was found in the analysis of motifs of all TFs (Additional file 1: Table S11). CTCF partially overlapped with cohesin RAD21 and SMC3 (Table 4). The top 20 pairs of FR and RF orientations of motifs overlapped or co-localized have been shown (Table 4). Unique pairs of overlapping or co-localized motifs were 404 FR and 662 RF, consisting of 117 FR and 218 RF of unique DNA motifs in monocytes. Only 10 overlapping or co-localized motifs were found to be common among the four cell types, including pairs of CTCF, RAD21, SMC3, and YY1, thereby implying that most overlapping and co-localized motifs serve in a cell-specific manner (Additional file 1: Table S11). Paired DNA-binding motifs of TFs with biased orientations tend to overlap, and a relatively small number of motifs are co-localized upstream and downstream of genes. Overlapping motifs act as TFBSs for several TFs either simultaneously or competitively [70–77]. CTCF and cohesin function cooperatively with another TF and a protein such as KLF4 and BRD2 [78, 79]. KLF4 recruits cohesin to the *Oct4* enhancer region. BRD2 supports boundary (insulator) activity of CTCF. The analysis of 457 ChIP-seq data sets from 119 human TFs showed that secondary motifs were detected in addition to the canonical motifs of the TFs, indicating tethered binding and co-binding between multiple TFs [80]. Wang et al. [80] observed significant position and orientation preferences between many co-binding TFs. Overlapping motifs found in this study may be biologically functional rather than artifact. Though repeat sequences in the human genome were masked, more than a hundred pairs of motifs overlapped with hundreds of upstream and downstream gene regions.

**Table 4.** Top 20 of the co-localized and overlapping biased orientations of DNA-binding motifs of TFs in monocytes. Co-localized and overlapping DNA motifs of CTCF with another biased orientation of DNA motifs are shown in separate tables. Motif 1 and Motif 2: DNA-binding motifs of TFs. No.: the number of genomic locations with both Motif 1 and Motif 2 upstream and downstream of genes in an EPA domain (< 1 Mb distance from TSS of genes). The total number of genes was 30,523. Co-localized and overlapping DNA motifs of CTCF with other DNA motifs.

Forward-reverse (FR)
| Motif 1 | Motif 2 | No. | Mean of distances bet. DNA sites (nt) | Median of distances bet. DNA sites (nt) | Mean of distances from TSS (nt) | Median of distances from TSS (nt) | Interquartile range of distance from TSS (nt) |
| --- | --- | --- | --- | --- | --- | --- | --- |
| FOS | JUN | 3332 | 0.00 | 0 | 43202.95 | 18085 | 5524 - 48583 |
| **CTCF** | **SMC3** | 2876 | 0.01 | 0 | 34606.92 | 13716 | 3073 - 39471 |
| FOSL2 | JUN | 2807 | 0.00 | 0 | 40389.84 | 17365 | 5424 - 45251 |
| FOSB | FOSL2 | 2644 | 0.00 | 0 | 44302.93 | 17867 | 5633 - 47209 |
| EGR1 | EGR2 | 2397 | 0.00 | 0 | 14578.21 | 1251 | 247 - 11473 |
| **CTCF** | **RAD21** | 2352 | 0.00 | 0 | 33492.68 | 12913 | 2824 - 37672 |
| **RAD21** | **SMC3** | 2343 | 0.02 | 0 | 33572.80 | 14302 | 3856 - 37159 |
| ATF3 | FOSL2 | 2322 | 0.00 | 0 | 42784.04 | 17841 | 5366 - 48236 |
| FOSL2 | JUNB | 2305 | 0.02 | 0 | 42884.36 | 17867 | 5706 - 46515 |
| ETS2 | ETSLIKE | 2205 | 0.00 | 0 | 37660.66 | 15765 | 4146 - 41462 |
| FOS | JUNB | 2151 | 0.00 | 0 | 38403.20 | 16367 | 4849 - 44736 |
| IRF | IRF4 | 1962 | 0.00 | 0 | 40794.41 | 15872 | 3969 - 45946 |
| FOSL2 | SMARCC1 | 1945 | 0.00 | 0 | 42429.69 | 17369 | 5094 - 46586 |
| EGR1 | WT1 | 1941 | 0.16 | 0 | 12485.23 | 427 | 93 - 5394 |
| FOSB | JUN | 1808 | 0.02 | 0 | 36648.74 | 16705 | 5156 - 40309 |
| ATF3 | JUN | 1713 | 0.02 | 0 | 38101.24 | 17119 | 5294 - 43559 |
| JUN | TFAP2A | 1628 | 0.00 | 0 | 42262.97 | 18017 | 5089 - 47094 |
| FOSL2 | TFAP2A | 1628 | 0.00 | 0 | 36606.38 | 16201 | 4949 - 41948 |
| BACH2 | SMARCC1 | 1462 | 0.16 | 0 | 38673.02 | 14778 | 4060 - 41262 |
| ELF1 | ETS1 | 1459 | 0.00 | 0 | 37056.72 | 14346 | 3370 - 40339 |

Reverse-forward (RF)
| Motif 1 | Motif 2 | No. | Mean of distances bet. DNA sites (nt) | Median of distances bet. DNA sites (nt) | Mean of distances from TSS (nt) | Median of distances from TSS (nt) | Interquartile range of distance from TSS (nt) |
| --- | --- | --- | --- | --- | --- | --- | --- |
| EGR1 | EGR3 | 3012 | 0.00 | 0 | 17005.00 | 709 | 149 - 10393 |
| NR1I2 | NR1I3 | 2932 | 0.00 | 0 | 49622.84 | 20548 | 5924 - 58436 |
| ZIC1 | ZIC3 | 2871 | 0.07 | 0 | 21511.64 | 3092 | 434 - 19128 |
| EGR | WT1 | 2612 | 0.04 | 0 | 9949.38 | 396 | 102 - 4266 |
| FOS | FOS:JUN | 2512 | 0.01 | 0 | 37200.80 | 15466 | 4438 - 42373 |
| EGR | EGR3 | 2293 | 0.17 | 0 | 11634.92 | 532 | 135 - 6044 |
| MAFG | NFE2L1:MAFG | 2272 | 0.03 | 0 | 34641.56 | 16917 | 5284 - 42097 |
| KLF10 | SP1 | 2233 | 0.00 | 0 | 9778.94 | 297 | 73 - 3077 |
| EGR | EGR2 | 2160 | 0.16 | 0 | 10502.76 | 490 | 129 - 5227 |
| KLF3 | SP3 | 2036 | 0.00 | 0 | 6974.78 | 246 | 63 - 1447 |
| KLF1 | SP3 | 1771 | 0.00 | 0 | 7748.42 | 264 | 66 - 1829 |
| KLF10 | SP3 | 1734 | 0.00 | 0 | 7275.48 | 222 | 58 - 1320 |
| KLF7 | SP4 | 1639 | 0.00 | 0 | 12521.62 | 309 | 58 - 4894 |
| KLF1 | SP1 | 1574 | 0.05 | 0 | 6780.46 | 245 | 61 - 1438 |
| AP1 | JUNB | 1572 | 0.05 | 0 | 34158.07 | 15451 | 4746 - 40315 |
| EGR1 | WT1 | 1543 | 0.31 | 0 | 12742.28 | 444 | 111 - 5553 |
| EGR1 | SP1:SP3 | 1393 | 0.08 | 0 | 7789.54 | 307 | 91 - 1886 |
| FOS:JUN | JUNB | 1386 | 0.14 | 0 | 33012.80 | 14245 | 3900 - 37988 |
| SP1 | SP3 | 1372 | 0.00 | 0 | 8845.31 | 293 | 77 - 1816 |
| EHF | ELF1 | 1366 | 0.02 | 0 | 41217.39 | 13357 | 2842 - 43178 |

Co-localized and overlapping DNA motifs of CTCF with other DNA motifs. FR orientation of CTCF
| Motif 1 | Motif 2 | No. | Mean of distances bet. DNA sites (nt) | Median of distances bet. DNA sites (nt) | Mean of distances from TSS (nt) | Median of distances from TSS (nt) | Interquartile range of distance from TSS (nt) |
| --- | --- | --- | --- | --- | --- | --- | --- |
| CTCF | SMC3 | 2876 | 0.01 | 0 | 34606.92 | 13716 | 3073 - 39471 |
| CTCF | RAD21 | 2352 | 0.00 | 0 | 33492.68 | 12913 | 2824 - 37672 |
| CTCF | CTCFL | 1151 | 0.00 | 0 | 30252.04 | 11606 | 2229 - 34539 |
| CTCF | ZBTB7A | 470 | 0.27 | 0 | 23227.96 | 7065 | 745 - 23496 |
| CTCF | E2F | 123 | 0.81 | 0 | 7913.36 | 222 | 61 - 1997 |
| CTCF | WT1 | 120 | 0.00 | 0 | 3543.72 | 114 | 44 - 375 |
| CTCF | MYC | 110 | 2.30 | 0 | 31289.45 | 11948 | 2887 - 33972 |

RF orientation of CTCF
| Motif 1 | Motif 2 | No. | Mean of distances bet. DNA sites (nt) | Median of distances bet. DNA sites (nt) | Mean of distances from TSS (nt) | Median of distances from TSS (nt) | Interquartile range of distance from TSS (nt) |
| --- | --- | --- | --- | --- | --- | --- | --- |
| CTCF | RXRA | 1362 | 0.00 | 0 | 29981.33 | 10937 | 2100 - 31407 |
| CTCF | CTCFL | 429 | 0.00 | 0 | 25171.22 | 9834 | 1859 - 24741 |
| CTCF | RAD21 | 337 | 0.00 | 0 | 38829.91 | 17973 | 6117 - 45749 |
| CTCF | ZIC1 | 126 | 1.40 | 0 | 35480.65 | 10286 | 2316 - 29380 |
| CTCF | ZIC3 | 108 | 0.00 | 0 | 30219.32 | 10904 | 4463 - 31037 |

## Discussion

A systematic method to find DNA-binding motifs of TFs that affect EPIs and gene expression without using chromatin interaction data is presented in this study. DNA motifs of TFs such as CTCF and cohesin (RAD21 and SMC3) were searched in the open chromatin regions of monocytes, T cells, HMEC, and NPC. A total of 96 biased (64 FR and 52 RF) orientations of DNA motifs were discovered and found to be common among the four cell types. Remarkably, non-oriented (i.e., without considering orientation) motifs was not found to affect the expression level of putative transcriptional target genes, suggesting that an EPA domain limited within the FR or RF orientation of TFBSs was a more accurate prediction of EPIs. This was supported by the comparisons with chromatin interaction and gene expression data. FF and RR orientations of motifs were not examined in this analysis since CTCF has FR or RF orientation of their TFBSs [3]. Among the 96 biased orientations of DNA motifs studied, 14 were reported to be associated with chromatin interactions, and many other TFs have chromatin-associated functions. The enrichment analysis of biased orientations of TFBSs of the 96 TFs for pairs of chromatin interaction anchors indicated a weak statistical significance, except for CTCF, RAD21, SMC3, ATB3, and ZBTB7A, suggesting that the TFs were difficult to find using DNA motifs of TFs alone [5].

Without considering the difference of orientations among cell types, 175 biased orientations were found in common among the four cell types. These numbers (96 and 175) are the numbers of unique TFs with biased orientations of DNA motifs and not the numbers of unique DNA motifs. Possible causes of the discrepancy are as follows: (1) A TF or an isoform of the same TF may bind to different DNA-binding motifs, according to cell types and/or in the same cell type. About 50% or more alternative splicing isoforms are differently expressed among tissues, thereby indicating that most alternative splicing is subject to tissue-specific regulation [81–83]; (2) The same TF has several DNA-binding motifs, and in some cases, one of the motifs is almost the same as the reverse complement motif of another motif of the same TF in a database; (3) I previously found that a complex of TFs would bind to a slightly different DNA-binding motif by combining DNA-binding motifs from the complex in *C. elegans* [84]. In other reports, the difference of binding specificity of TF pairs was investigated [85]. Cofactors also alter the binding specificity of TFs [70–72]. These increase the variety and number of (biased orientations of) DNA-binding motifs, which are potentially involved in different biological functions among cell types.

To predict TFBSs in genomic DNA, several tools and methods were explored and then protein interaction quantitation (PIQ) was used [27]. PIQ predicts TFBSs by reducing noise in DNase-seq data, and the comparison of the prediction with ChIP-seq data has revealed that PIQ shows a better performance than other tools and digital genome footprinting (DNase-DGF). The prediction of TFBSs using PIQ in this study increased the number of biased orientations of motifs, thereby reducing the number of non-oriented motifs. At first, the number of predicted motifs was expected to increase relative to using consensus DNA-binding sequences of TFs and open chromatin regions available in the ENCODE database, which seemed to be difficult to find all TFBSs in the genome. However, the number of TFBSs was almost the same or slightly decreased compared with the analysis of consensus sequences, suggesting that PIQ increased true-positive predictions of TFBSs and reduced false-positive predictions.

When forming a homodimer or heterodimer with another TF, a TF may bind to genomic DNA with a specific orientation of its TFBSs. The approach in this study would be able to identify TFs forming a heterodimer. If the TFBS of only the mate to a pair of TFs was found in an EPA domain, the EPA domain was limited within one side of the TFBS, and transcriptional target genes were predicted using the EPA domain limited within the side. In this analysis, the mate to both heterodimers and homodimers of TFs can be tested to examine the effect on the expression level of transcriptional target genes predicted using the EPA domain limited within one side. Biased orientations of DNA-binding motifs may also be found in forward-forward or reverse-reverse orientation.

TFs that are multi-functional transcriptional regulators such as YY1 may not indicate a strong directional bias of their TFBSs. Though some proportion of TFBSs of a TF involved in chromatin interactions and insulator functions is found in a biased orientation in the genome, the other TFBSs of the TF may not form a dimer with another TF and may not be present in a specific orientation in the genome. Thus, the overall proportion of biased orientations of TFBSs of a TF may be reduced such that a statistical test cannot detect it.

The approach presented in this study was tested to search for repeat sequences that serve as insulator sites and affect the expression level of putative transcriptional target genes of TFs. A small number of repeat sequences were identified and included MIR, which act as insulators [86].

CTCF and cohesin-binding sites are frequently mutated in cancer [87]. Some biased orientations of DNA motifs involved in chromatin interactions may be associated with diseases including cancer.

The analyses conducted here suggest that biased orientations of DNA motifs of discovered TFs affect EPIs and the expression level of putative transcriptional target genes of general TFs bound in enhancers and promoters. This gives insight into the nature of TFs and contributes to the understanding of functions and molecular mechanisms of chromatin interactions, regulation of chromatin, histone modifications, transcriptional regulation, and transcriptional cascades.

## Methods

### Search for biased orientations of DNA motifs

To identify transcription factor binding sites (TFBSs) from open and closed chromatin regions, TRANSFAC (2019.2), JASPAR (2018), UniPROBE (2018), high-throughput SELEX, transcription factor binding sequences of ENCODE ChIP-seq data, and HOCOMOCO version 9 and 11 were used to predict insulator sites [88–94]. TRANSFAC (2019.2) was used to analyze enhancer-promoter interactions because these data were sufficient to identify biased orientations of DNA motifs of insulator TFs with less computational time, thereby reducing the number of non-oriented DNA motifs of TFs. To reduce false-positive predictions of TFBSs in open chromatin regions, the DNA motif search was performed by using the protein interaction quantitation (PIQ) tool with 12,249 position frequency matrices converted from DNA motifs of vertebrate TFs in the above-mentioned databases and DNase-seq data from GEO and ENCODE database (GSM1024791 CD14^+^ monocytes; GSM665812 CD4^+^ T cell; GSM736634 Human mammary epithelial cell, HMEC; GSM878615 Neural progenitor cell, NPC; ENCFF263GMV, ENCFF886XEV Th17; GSM1024741 Treg; GSM736620 GM12878) [27]. To find open chromatin regions associated with enhancer and promoter activity, narrow peaks of ChIP-seq experimental data of H3K27ac were predicted using MACS2 callpeak (GEO: GSM773004) [95]. To predict TFBSs of TFs from closed chromatin regions, another method was employed. Position weight matrices of vertebrate transcription factor binding sequences were converted into DNA consensus sequences, which consist of an alphabet of 15 characters (the four bases A, C, G, T, the six two-fold degenerate IUPAC codes R=[AG], Y=[CT], K=[GT], M=[AC], S=[GC], W=[AT], the four three-fold degenerate IUPAC codes B=[CGT], D=[AGT], H=[ACT], V=[ACG], and the four-fold degenerate character N=[ATGC]) using a convert matrix in the RSAT website [96]. Transcription factor binding sequences were searched from genomic regions other than narrow peaks of DNase-seq data in repeat-masked hg19 genome sequences using the Match tool in TRANSFAC with a similarity score cutoff of 1 after transforming the DNA consensus sequences of TFs into position frequency matrixes for the Match search [97]. Repeat-masked hg19 genome sequences were downloaded from the UCSC genome browser (http://genome.ucsc.edu/, http://hgdownload.soe.ucsc.edu/goldenPath/hg19/bigZips/hg19.fa.masked.gz). TFs corresponding to transcription factor binding sequences were searched computationally and manually by comparing their names and gene symbols of HUGO Gene Nomenclature Committee-approved gene nomenclature and 31,848 UCSC canonical transcripts (http://hgdownload.soe.ucsc.edu/goldenPath/hg19/database/knownCanonical.txt.gz) because transcription factor binding sequences were not linked to transcript IDs such as UCSC, RefSeq, and Ensembl transcripts.

Target genes of a TF were assigned when its TFBS was found in the promoter or extended regions of a gene’s EPA. Promoter and extended regions were defined as follows: promoter regions were those that were within a distance of ±5 kb from transcriptional start sites (TSS). Promoter and extended regions were defined as per the following association rule, which is the same as that defined in Figure 3A of a previous study [12]: the single nearest gene association rule, which extends the regulatory domain to the midpoint between the TSS of the gene and that of the nearest gene upstream and downstream without the limitation of extension length. The limitation of extension length was <1 Mb in this study. Extended regions for an EPA were limited within the TFBSs of a putative insulator TF closest to a TSS, and transcriptional target genes were predicted using the limited EPA domain and TFBSs of a general TF in open chromatin regions. Promoter and extended regions of an EPA were limited within the forward–reverse (FR) orientation of TFBSs of a putative insulator TF. When the forward or reverse orientation of TFBSs was located in the genome several times, the most internal (i.e., closest to a TSS) forward–reverse orientation of the TFBSs was selected. The genomic positions of genes were identified using the ‘knownGene.txt.gz’ file in UCSC bioinformatics sites [98]. The file ‘knownCanonical.txt.gz’ was also utilized to select representative transcripts among various alternate forms for assigning promoter and extended regions of an EPA. From the list of TFBSs and transcriptional target genes, redundant DNA motifs of TFs were removed by comparing the putative target genes of a TFBS and its corresponding TF; if identical, one of the DNA motifs of TF was chosen. When the number of transcriptional target genes predicted based on a TFBS was <5, the DNA motif of TF was omitted.

For gene expression data, RNA-seq reads mapped onto the human hg19 genome sequence were obtained from UCSF-UBC human reference epigenome mapping project RNA-seq reads with poly-A of naive CD4^+^ T cells (GEO: GSM669617). For monocytes, Blueprint RNA-seq FPKM data (‘C0010KB1.transcript_quantification.rsem_grape2_crg.GRCh38.20150622.results’) were downloaded from Blueprint DCC portal (http://dcc.blueprint-epigenome.eu/#/files). RNA-seq reads with the poly-A of human mammary epithelial cells (HMEC) in the Encyclopedia of DNA Elements at UCSC (http://genome.ucsc.edu/encode/, ‘wgEncodeCshlLongRnaSeqHmecCellPapAlnRep1.bam’ file), H1-derived NPC (GEO: GSM915326, ENCODE: ENCFF529SIO), Th17 (GSM2859479, NCBI SRA: SRR6298326), Treg (GSM2859476, SRR6298323), and GM12878 (GSE78551, ENCFF297QCE) were analyzed. RNA-seq reads were aligned in repeat-masked hg19 genome sequences using Burrows-Wheeler Aligner with default parameters [99]. FPKMs of the RNA-seq data were calculated by RSeQC [100]. Based on log2-transformed FPKM, UCSC transcripts were arranged in a descending order of expression level, and the top 4,000 transcripts were selected in each cell type.

The expression level of transcriptional target genes predicted using an EPA domain limited within the TFBSs of a TF was compared with the expression level of transcriptional target genes predicted using closed chromatin regions in promoters. For each DNA motif limiting the EPA domain, transcriptional target genes were predicted based on DNA-binding motifs of vertebrate TFs in the TRANSFAC database, and the ratio of expression level of putative transcriptional target genes of each TF was calculated between the EPA domain and closed chromatin regions in the promoters. The distribution of the ratios of all TFs was compared among forward-reverse (FR), reverse-forward (RF), and non-orientation (i.e., without considering orientation) of TFBSs limiting an EPA domain using a two-sided Mann-Whitney test, (*p*-value < 10^−7^). Other parameters for the analysis were set as follows: the number of transcriptional target genes of a TF was ≥50, the number of general TFs (not insulator TFs) to predict transcriptional target genes was ≥50, and the number of TFBSs of a TF (insulator TF) shortening an EPA was ≥100.

### Comparison with chromatin interaction data

For comparison of EPIs with chromatin interactions (HiChIP) in CD4^+^ T, Th17, Treg, and GM12878 cells, ‘GSM2705049_Naive_HiChIP_H3K27ac_B2T1_allValidPairs.txt’, ‘GSM2705050_Naive_HiChIP_H3K27ac_B2T2_allValidPairs.txt’, ‘GSM2705051_Naive_HiChIP_H3K27ac_B3T1_allValidPairs.txt’, ‘GSM2705054_Th17_HiChIP_H3K27ac_B1T2_allValidPairs.txt’, ‘GSM2705055_Th17_HiChIP_H3K27ac_B2T1_allValidPairs.txt’, ‘GSM2705056_Th17_HiChIP_H3K27ac_B3T1_allValidPairs.txt’, ‘GSM2705059_Treg_HiChIP_H3K27ac_B3T1_allValidPairs.txt’, ‘GSM2705041_GM_HiChIP_H3K27ac_B1_allValidPairs.txt’, and ‘GSM2705042_GM_HiChIP_H3K27ac_B2_allValidPairs.txt’ files were downloaded from the Gene Expression Omnibus (GEO) database. The resolutions of chromatin interactions and EPIs were adjusted to 5 kb before comparison. Chromatin interactions with ≥ 6,000 and ≥ 1,000 counts for each interaction were analyzed in the comparison in CD4^+^ T cells. According to the unique numbers of chromatin interactions in replications in Th17, Treg, and GM12878 cells, chromatin interactions with ≥1,000 (B1T2 in Th17), ≥2,000 (B2T1 and B3T1 in Th17), ≥ 6,000 (B3T1 in Treg), and ≥4,000 (B1 and B2 in GM12878) counts for each interaction were analyzed in the comparison.

EPIs were predicted on the basis of the following three types of EPA domains in monocytes: (i) EPA domain limited within the FR or RF orientation of TFBSs of a TF, EPA domain limited within the non-oriented (i.e., without considering orientation) TFBSs of a TF, and (iii) EPA domain without being limited within a TFBS. EPIs predicted using the three types of EPA domains in common were removed. EPIs predicted using EPA domain (i) were removed from EPIs predicted using EPA domain (ii). EPIs predicted using EPA domain (i) and (ii) were removed from EPIs predicted using EPA domain (iii). The resolution of HiChIP chromatin interaction data was 1-5 kb; thus, EPIs were adjusted to 5 kb before comparison. The number and ratio of EPIs overlapped with chromatin interactions were compared two times between EPI (i) and (iii), and EPI (i) and (ii) (binomial distribution, *p*-value < 0.025 for each test, two-sided, 95% confidence interval).

For comparison of EPIs with chromatin interactions (in situ Hi-C) in HMEC, the genomic positions of chromatin interactions in the ‘HMEC_4DNFI97O9SAZ.pairs.gz’ file in 4D nucleome data portal (http://data.4dnucleome.org) were converted into the hg19 version of human genome using the liftOver tool (http://hgdownload.soe.ucsc.edu/downloads.html#liftover). The resolutions of chromatin interaction data and EPIs were adjusted to 3 kb before comparison. All chromatin interactions were used in the comparison. For comparison of EPIs with chromatin interactions (promoter capture Hi-C) in NPC, the resolutions of chromatin interaction data in the ‘GSE86189_npc.po.all.txt.bz2’ file (GEO: GSE86189) and EPIs were adjusted to 5 kb before comparison. All chromatin interactions were used in the comparison.

Putative target genes for the comparison of EPIs and chromatin interactions were selected from top 4,000 transcripts in terms of the expression level. The expression level of putative target genes of EPIs overlapped with chromatin interactions was compared with EPIs not overlapped with them. When the same transcriptional target gene of a TF in an enhancer was predicted using both an EPI overlapping a chromatin interaction and a non-overlapping EPI, the target gene was eliminated. The distribution of expression level of putative target genes was compared using a two-sided Mann-Whitney test (*p*-value < 0.05).

Micro-C chromatin interaction data in human fibroblasts (4D nucleome data portal ID: 4DNFICOEXGPJ) was converted into the hg19 version of the human genome using the liftOver tool and was adjusted to 200-bp resolution. The same genomic locations of paired chromatin interaction anchors were merged.

### Correlation of expression level of gene pairs separated by TFBS

To examine the correlation of expression level of genes pairs separated by TFBSs of a TF, gene expression data among 53 tissues were obtained from the ‘GTEx_Analysis_2016-01-15_v7_RNASeQCv1.1.8_gene_median_tpm.gct.gz’ file in the GTEx database (https://gtexportal.org/home/). The correlation of log2-transformed gene expression levels was analyzed based on RefSeq protein-coding transcripts (RefSeq ID with ‘NM_’). The same genomic locations of RefSeq transcripts were removed, and TFBSs overlapped with RefSeq transcripts were also eliminated. Genes closest to a TFBS and another gene were analyzed using BEDOPS closest-features [101]. The Mann-Whitney test was performed using R.

## Abbreviations

TF: Transcription factors
TFBS: Transcription factor binding sites
TSS: Transcriptional start sites
EPA domain: Promoter and extended regions for enhancer-promoter association
EPI: Enhancer-promoter interaction
FF: Forward-forward
FR: Forward-reverse
RF: Reverse-forward
RR: Reverse-reverse
ChIP-seq: ChIP-sequencing, chromatin immunoprecipitation followed by massively parallel DNA sequencing
DNase-seq: DNase I hypersensitive sites sequencing
RNA-seq: RNA sequencing
ENCODE: Encyclopedia of DNA elements

## Acknowledgements

Supercomputing resources were provided by the Human Genome Center of the Institute of Medical Science at the University of Tokyo. This work was partially supported by JSPS KAKENHI (Grant Number 16K00387) and JST CREST (Grant Number JPMJCR15G1), Japan.

