## Supplemental data 2 for "Discovery of directional chromatin-associated regulatory motifs affecting human gene transcription"

### Results

Additionally, to confirm the tendencies of comparison of EPIs with HiChIP chromatin interactions, the same analysis was conducted using HiChIP chromatin interaction data in Th17, Treg, and GM12878 cells. For a total of 128 biased orientations [76 of 200 (38%) FR and 64 of 183 (35%) RF] of DNA motifs in Th17 cells, which included CTCF, cohesin (RAD21 and SMC3), ZNF143, and YY1 in three replications (B1T2, B2T1, and B3T1), a significantly higher ratio of EPIs overlapped with HiChIP chromatin interactions in EPA domain (i) than in EPA domain (ii) and (iii) (Additional file 1: Table S6). When comparing EPIs predicted using only EPA domain (i) and (iii) with the chromatin interactions, for a total of 282 biased orientations [182 of 200 (91%) FR and 162 of 183 (89%) RF] of DNA motifs, a significantly higher ratio of EPIs overlapped with the chromatin interactions in EPA domain (i) than EPA domain (iii) (Additional file 1: Table S6).

In Treg cells, chromatin interactions in one biological replication were overlapped with EPIs, but because those in the other two biological replications were less overlapped with EPIs, one replication (B3T1) was used for this analysis. For a total of 281 biased orientations [154 of 290 (53%) FR and 176 of 323 (54%) RF] of DNA motif sequences, which included CTCF, cohesin (RAD21 and SMC3), ZNF143, and YY1, a significantly higher ratio of EPIs overlapped with HiChIP chromatin interactions in EPA domain (i) than the other types of EPA domain (ii) and (iii) (Additional file 1: Table S6). When comparing EPIs predicted using only EPA domain (i) and (iii) with the chromatin interactions, for total 482 biased orientations [283 of 290 (98%) FR and 309 of 323 (96%)

RF] of DNA motif sequences, a significantly higher ratio of EPIs overlapped with the chromatin interactions in EPA domain (i) than EPA domain (iii) (Additional file 1: Table S6).

In GM12878 cells, for a total of 168 biased orientations [96 of 229 (38%) FR and 98 of 237 (35%) RF] of DNA motifs including CTCF, cohesin (RAD21 and SMC3), ZNF143, and YY1 in two replications (B1 and B2), a significantly higher ratio of EPIs overlapped with HiChIP chromatin interactions in EPA domain (i) than EPA domain (ii) and (iii) (Additional file 1: Table S6). When comparing EPIs predicted using only EPA domain (i) and (iii) with the chromatin interactions, for a total of 366 biased orientations [218 of 229 (95%) FR and 227 of 237 (96%) RF] of DNA motifs, a significantly higher ratio of EPIs overlapped with the chromatin interactions in EPA domain (i) than EPA domain (iii) (Additional file 1: Table S6).
